# Genetic determinants in *Salmonella enterica* serotype Typhimurium required for overcoming stressors in the host environment

**DOI:** 10.1101/571273

**Authors:** Rabindra K. Mandal, Tieshan Jiang, Young Min Kwon

**Affiliations:** Department of Poultry Science, University of Arkansas, Fayetteville, AR, 72701; Cell and Molecular Biology Program, University of Arkansas, Fayetteville, AR, 72701

**Keywords:** *Salmonella*, host stress, Tn-seq, conditionally essential genes, *in vitro* fitness

## Abstract

*Salmonella enterica* serovar Typhimurium (S. Typhimurium), a non-typhoidal *Salmonella* (NTS), result in a range of diseases, including self-limiting gastroenteritis, bacteremia, enteric fever, and focal infections representing a major disease burden worldwide. There is still a significant portion of *Salmonella* genes whose functional basis to overcome host innate defense mechanisms, consequently causing disease in host, largely remains unknown. Here, we have applied a high-throughput transposon sequencing (Tn-seq) method to unveil the genetic factors required for the growth or survival of S. Typhimurium under various host stressors simulated *in vitro*. A highly saturating Tn5 library of *S*. Typhimurium 14028s was subjected to selection during growth in the presence of short chain fatty acid (100 mM propionate), osmotic stress (3% NaCl) or oxidative stress (1 mM H_2_O_2_) or survival in extreme acidic pH (30 min in pH3) or starvation (12 days in 1X PBS). We have identified an overlapping set of 339 conditionally essential genes (CEGs) required by *S*. Typhimurium to overcome these host insults. Interestingly, entire eight genes encoding F_0_F_1_-ATP synthase subunit proteins were required for fitness in all five stresses. Intriguingly, total 88 genes in *Salmonella* pathogenicity island (SPI), including SPI-1, SPI-2, SPI-3, SPI-5, SPI-6 and SPI-11 are also required for fitness under the *in vitro* conditions evaluated in this study. Additionally, by comparative analysis of the genes identified in this study and the genes previously shown to be required for *in vivo* fitness, we identified novel genes (*marBCT*, *envF*, *barA*, *hscA*, *rfaQ*, *rfbI* and putative proteins STM14_1138, STM14_3334, STM14_4825, and STM_5184) that has compelling potential to be exploited as vaccine development and/or drug target to curb the *Salmonella* infection.

## Introduction

Non-typhoidal *Salmonella* (NTS), a Gram-negative bacterial pathogen, causes 93 million enteric infections, 155,000 diarrheal deaths, and 3.4 million blood stream infection worldwide annually (Ao et al., 2015; Majowicz et al., 2010). Gram-negative bacterial pathogens, including NTS, are developing resistance against antimicrobial agents including the last resort antibiotics at a startling rate, creating a global crisis in human health. Scientists fear the impending global epidemic of untreatable infections and return to a pre-antibiotic era where a common infection and minor injury can be lethal (Liu et al., 2015; McKenna, 2013; Spencer, 2015; World Health Organization (WHO). Thus, there is an urgent need to identify genetic factors of pathogenic microorganisms that can serve as targets to develop novel strategies to combat infectious diseases (Medini et al., 2008; van Opijnen and Camilli, 2012). Nonetheless, the insufficiency of the genome-wide data that provide links between genotype and the infection-related phenotypes of bacteria is the major roadblock to discover suitable targets for development of the effective strategies to control infection.

*Salmonella enterica* serotype Typhimurium (*S.* Typhimurium) is one of the leading cause of NTS (Carden et al., 2015; Crim et al., 2015). Despite *Salmonella* infection has an enormous global burden on disease worldwide and availability of complete genome sequence of *S.* Typhimurium LT2 nearly one and half decade (2002) ago, the phenotypic basis of *S.* Typhimurium genes required for *in vivo* survival is still unknown for a large portion of the genes (Feasey et al., 2012; McClelland et al., 2001). Researchers have tried to delve into the pathogenesis of *S.* Typhimurium using different variations of high throughput screening of transposon mutants, with a limited number of mutants based on a negative selection (Kwon et al., 2016). Chan et al., (2005) had discovered 157 and 264 genes required by *S.* Typhimurium strain SL1344 for acute infection in mice (A-Mice) and survival inside macrophage (MΦ), respectively using a microarray-based tracking method (Chan et al., 2005). Lawley et al., (2006) used the same method to identify 118 genes of *S.* Typhimurium SL1344 required for long-term persistent infection in mice (P-Mice) using the spleen samples collected after 28 day post infection (Lawley et al., 2006). Additionally, Chaudhuri et al. (2013) have comprehensively assigned a core set of 611 genes of *S.* Typhimurium strain ST4/74 required for effective colonization in the calf, pig, and chicken (Chaudhuri et al., 2013). Recently, Silva-Valenzuela et al., identified 224 mutants of *S*. Typhimurium 14028s that were negatively selected using two pools of single gene deletion mutants from spleen and liver at 2 days post infection in mice (Sp-Liv) (Silva-Valenzuela et al., 2015). Previously, our laboratory conducted Tn-seq screening to identify an overlapping set of 105 coding genes of *S*. Typhimurium 14028s required for *in vitro* growth in diluted Luria-Bertani (LB) medium, LB medium plus bile acid and LB medium at 42°C (Khatiwara et al., 2012). However, there is still a gap in the above approaches to correlate *in vivo* and *in vitro* survival or growth genes required by *S.* Typhimurium that will help delve into biochemical and molecular basis of virulence and potentially pave a roadmap towards the efficient development of novel vaccines, antibiotics, and control strategies.

In this study, we conducted transposon sequencing (Tn-seq) analysis of *S.* Typhimurium 14028s under the five *in vitro* conditions mimicking host stressors found during enteric and systemic infection. Tn-seq is a powerful tool for functional analysis of bacterial genomes based on the use of random transposon mutagenesis and next generation sequencing technology (Kwon et al., 2016; Van Opijnen et al., 2009; Van Opijnen and Camilli, 2013). We have applied a highly efficient method for Tn-seq library preparation that requires only small amount of DNA without the need for enzymatic digestion or physical shearing of genomic DNA (Dawoud et al., 2014; Karash et al., 2017; Mandal and Kwon, 2017; Mandal et al., 2017). To cause enteric infection *S.* Typhimurium has to overcome gastrointestinal host insult such as low acidic pH in the stomach, osmotic and short chain fatty acid (SCFAs) in intestine (Ha et al., 1998; Nava et al., 2005; Sleator and Hill, 2002; Smith, 2003). Eventually, for systemic infection, *S.* Typhimurium has to vanquish macrophage stress such as oxidative stress, starvation as well as hyperosmotic condition (Lee et al., 2014; Rosenberger and Finlay, 2003; van der Heijden et al., 2015). We hypothesized that the comparative analysis of the comprehensive sets of the *in vitro* fitness genes (for stress resistance, this study and previous) and *in vivo* (required for enteric and systemic infection in the host) will allow better understanding of the biochemical or phenotypic basis of the genetic requirements of *S.* Typhimurium for host infection and provide enhanced resolution to link genotype to phenotype. Thus, we performed a comparative study between the *in vivo* and *in vitro* fitness genes from previous studies and this study, respectively.

## Material and Methods

### Bacterial strains and growth conditions

*S.* Typhimurium 14028s, a spontaneous mutant resistant to nalidixic acid (NA), was grown in Luria-Bertani (LB) plate or LB medium (BD Difco, Sparks, MD) on shaking rack at 225 rpm and incubated at 37°C unless otherwise indicated. Nalidixic acid (NA, ICN Biomedicals Inc., Aurora OH, USA) and Kanamycin (Km, Shelton Scientific, Inc. CT, USA) were used at 25 μg/ml and 50 μg/ml respectively. *S.* Typhimurium was stored in 50% glycerol at −80°C.

### Construction of Transposon mutant library

To prepare electrocompetent cells, *S.* Typhimurium was grown overnight in 10 ml LB medium with NA and was diluted 100 fold in 10 ml 2xYT (BD Difco, Sparks, MD, USA) medium with NA and incubated for 3 h on a shaking rack. Bacterial cells were washed 6 times with wash solution (10% glycerol). Centrifugation was done at 8,000 rpm for 1 min at refrigeration temperature (4°C). The bacterial pellet was mixed gently in 60 μl of wash solution preventing aeration. One μl of the EZ-Tn5 <KAN-2> Tnp transposome complex (Epicentre BioTechnologies, Madison, WI, USA) was added to electrocompetent *S.* Typhimurium cells and incubated on ice for 10 min. Then, the mixture was gently transferred to ice cold cuvette avoiding the formation of any air bubble and electroporated at 2450 V. Immediately, 500 μl of SOC was added and incubated for 90 minutes on a shaking rack at 37°C. The reaction was plated on LB plates supplemented with NA and Km to recover the transformants. With three electroporations we were able to collect 350,000 Tn5 mutants and stored them in LB medium with 50% glycerol at −80°C (Figure 1).

**Figure 1.**
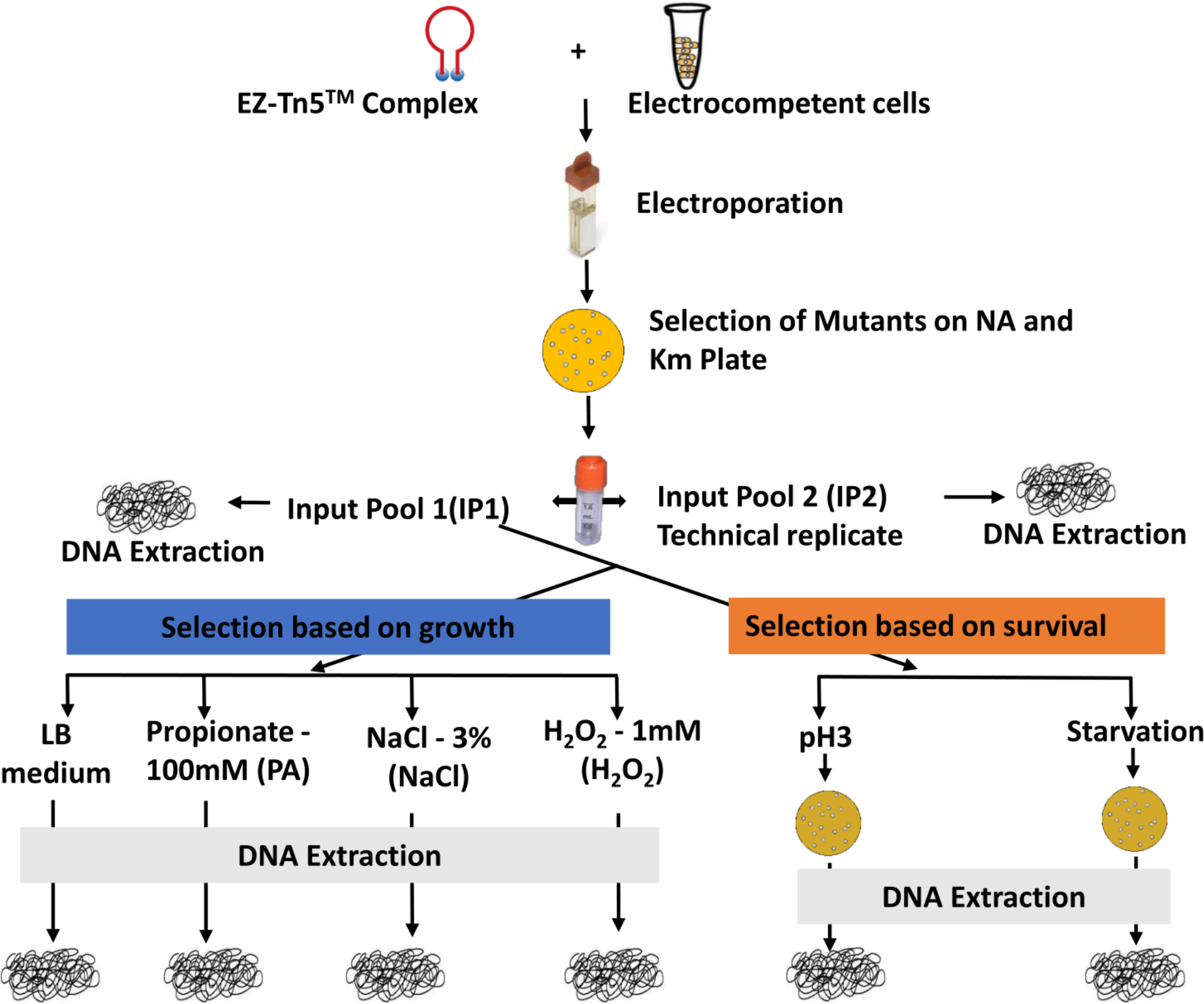
Schematic overview of the experimental design. A highly saturating Tn5 library was constructed through electroporation of EZ-Tn5 transposome complex to *S*. Typhimurium 14028s. Approximately 350,000 Tn5 mutants were collected on LB (Km + NA) plates. Complex Tn5 mutant library (IP1) was selected based on growth [LB medium (LB), 100 mM Propionate in LB medium (PA), 3% NaCl in LB medium (NaCl), and 1mM Hydrogen peroxide in LB medium (H_2_O_2_)] and survival [exposed to pH3 for 30 min (PH3) and incubated for 12 days in 1X PBS (Starvation)]. Input pool 2 (IP2) was a technical replicate of input pool 1 (IP1).

### *In vitro* growth assay of transposon mutant library

*In vitro* selection of transposon mutant library was done as described by Opijnen and Camilli, (2010) (van Opijnen et al., 2014) with some modifications. Briefly, transposon mutant library was thawed on ice and an aliquot of 300 μl was added to 60 ml LB broth with NA and Km (OD_600_ = 0.131). The library was incubated at 37°C on a shaking rack for 30 min (OD_600_ = 0.135) and centrifuged at 5,500 rpm for 8 min at room temperature. The transposon mutant library pellet was resuspended in 50 ml 1X phosphate buffer saline (PBS) (OD_600_ = 0.143) and CFU (4×10^7^/ml) was measured (*t*_1_). This step was included to prepare the mutant cells adapted to LB medium and shorten the lag phase in the following selective conditions. Ten ml aliquot were saved from *t*_1_ as an input pool (IP1). Above procedure was repeated to make a technical replicate of IP1 as input pool 2 (IP2). An aliquot of 0.5 ml from *t*_1_ was inoculated to 10 ml LB (LB), LB with 3% NaCl (NaCl), LB with 100mM propionate with pH adjusted to pH7 (PA), LB with 1mM H_2_O_2_ (H_2_O_2_). The initial OD_600_ of inoculated medium was 0.009. We then incubated the libraries on a shaking rack (225 rpm) at 37°C with variable incubation time ranging from 3.75 h to 7 h (*t*_2_) to a mid-logarithmic. The final OD_600_ of all output pools was very similar around 0.64 at time point *t*_2_. Input pool and output pool libraries were centrifuged and the pellet was stored at −80°C for DNA extraction (Figure 1).

### *In vitro* survival assay of transposon mutant library

To identify genes negatively selected during starvation, an aliquot of 0.5 ml from t_1_ was transferred to 10 ml PBS and incubated at 37°C on shaking rack for 12 days. On the 12^th^ day, the tube was centrifuged and the pellet was dissolved in 1 ml PBS. 100 μl aliquot was incubated on LB plate (NA + Km) overnight at 37°C. The cells were collected in PBS and stored at −80°C for DNA extraction. Whereas for survival in pH3, 0.5 ml from *t*_1_ was exposed to LB medium adjusted at pH3 for 30 min at 37°C and immediately transferred to 40 ml PBS. The cells were centrifuged at 8000 rpm for 8 min and pellet was mixed in 1ml PBS. An aliquot of 250 μl was plated on LB plate (NA + Km) overnight at 37°C. Colonies were collected in PBS and stored at −80°C for DNA extraction (Figure 1).

### DNA library preparation for Illumina sequencing

Genomic DNA (gDNA) from the bacterial cell pellet of input library and output libraries stored at −80°C was extracted using QIAamp DNA Mini Kit (Qiagen, Valencia, CA, USA) following manufacturer’s protocol. The purity and concentration were checked using Qubit 2.0 Fluorometer (Life Technologies, Carlsbad, CA) with Qubit Assay Kits (dsDNA BR Assay) following the manufacturer’s manual.

The sample for Illumina sequencing was prepared as previously described (Dawoud et al., 2014; Mandal and Kwon, 2017; Mandal et al., 2017; Mandal, 2016). All the DNA primers (Table S5) used for Tn-seq library were custom designed using Primer3 (v. 0.4.0) (Untergasser et al., 2012) and ordered from Integrated DNA Technologies (Coralville, Iowa). The simplified diagram for preparation of Tn-seq amplicon library is shown Figure S1A. Briefly, Tn5-junctions at the right end of transposon was enriched from gDNA extracted from input and output library. The single primer linear extension was done with EZ-Tn5 primer3 using Taq DNA polymerase (New England Biolabs, Ipswich, MA, USA). The 50 μl linear PCR extension reaction constituted: Nuclease-free water – 40 μl (volume adjusted according to gDNA volume), Thermopol buffer (10X) – 5 μl, dNTPs (2.5 mM each) – 1 μl, EZ-Tn5 primer3 (20 μM) – 1 μl, gDNA library (50 ng/ul) – 2 μl (~100 ng), and Taq DNA polymerase – 1 μl (added during PCR). The PCR cycle consisted of manual hot start with the initial denaturation at 95°C for 2 min, and addition of Taq DNA polymerase followed by 50 cycles of 95°C for 30 s, 62°C for 45 s, and 72°C for 10 s, which was then followed by a hold at 4°C. The linear PCR products were then purified with MinElute PCR purification kit (Qiagen, Valencia, CA, USA) and eluted in 10 μl of elution buffer (EB) following the manufacturer’s protocol. Then deoxycytosine homopolymer tail (C-tail) was added to the linear extension purified PCR product using Terminal Transferase (TdT, New England Biolabs, Ipswich, MA, USA) enzyme following previous protocol (Lazinski and Camilli, 2013). The C-tailing reaction consisted: DNA (linear extension product from linear PCR) – 10 μl, TdT Buffer (10X) – 2 μl, CoCl_2_ (2.5 mM) – 2 μl, dCTP (10 mM) – 2.4 μl, ddCTP (1mM) – 1 μl, Nuclease-free H_2_O – 1.6 μl, and Terminal Transferase – 1 μl, making a total volume of 20 μl. The reaction was incubated at 37°C for 1 h followed by heat inactivation of the enzyme at 75°C for 20 min on a thermocycler. The C-tailed products were purified using MinElute PCR purification kit and eluted to 10 μl.

Subsequently, C-tailed PCR product was enriched with exponential PCR. PCR reaction constituted: nuclease-free H_2_O – 35 μl, Thermopol Buffer (10X) – 5 μl, dNTPs (2.5 mM each) – 4 μl, IR2 BC primer (with Illumina adapter and barcode, 10 μM) – 2 μl, HTM primer (with Illumina adapter, 20 μM) – 1 μl, C-tailed DNA – 2 μl, and Taq DNA Polymerase (NEB) – 1 μl, making a total volume of 50 μl. The manual hot start PCR cycle comprised of 95°C for 2 min, followed by 25 cycles of 95°C for 30s, 58°C for 45s, and 72°C for 20s, trailed by a final extension at 72°C for 10 min.

Finally, the exponential PCR products were pulse heated at 65°C for 15 min and ran on 1.5% agarose gel. Tn-seq library had smear pattern whereas gDNA of *S*. Typhimurium (negative control) had almost no amplification (Figure S1B). Gel was excised ranging from 300-500 bp and DNA was extracted using QIAquick Gel Extraction Kit (Qiagen, Valencia, CA). The purity and concentration of DNA were measured using Qubit 2.0 Fluorometer. An equal amount (~ 10 ng) of DNA (gel-purified products) from each library were mixed together and sent for next generation sequencing, Illumina HiSeq 2000 single end read 100 cycles (Center for Genome Research and Biocomputing, Oregon State University, Corvallis).

### Analysis of Transposon sequencing data

Raw reads from HiSeq Illumina sequencing were de-multiplexed based on the barcodes to their respective libraries using custom Perl script. The barcode and transposon sequence were trimmed off from 5’ end. Consequently, the remaining sequence was Tn5-junction sequences with/without poly C-tail. Only 20 bp from the Tn5-junction were kept discarding most of the poly C-tails. The reads were then aligned against *S.* Typhimurium 14028s complete genome (NC_016856.1) using Bowtie version 0.12.7(Langmead and Salzberg, 2012). The aligned sequence (SAM mapping file) were fed to ARTIST pipeline to identify conditionally essential genes (CEGs) using Con-ARTIST (Pritchard et al., 2014). Briefly, Tn5 insertion frequency was assigned to the *S.* Typhimurium 14028s genome divided into 100 bp window size. Uncorrected raw data (non-normalized) of input and output libraries were used to normalize the control data (IP1) to account for the random loss of mutants in output pool. Then, reads were compared between the matching input and output pool using a Mann-Whitney U test (MWU). The MWU results were used train hidden Markov model (HMM) to predict the likelihood of loci that were not required for growth in either condition, essential under both conditions, enriched in output library and window depleted in output library (p < 0.01). The insertions were only considered in the central 80% of the gene to avoid any polar effect of transposon insertion. The cutoffs for depleted loci and enriched loci were >8 fold and >2 fold, respectively.

### Comparative analysis of conditionally essential genes (CEGs) between *in vitro* and *in vivo* stressors

We compared the *in vitro* essential genes identified in this study and our previous study (Khatiwara et al. 2012) with the previously identified *in vivo* fitness genes. CEGs for acute infection of mice (A-Mice), macrophage survival (MΦ) (Chan et al., 2005) and persistent infection of mice (P-Mice) (Lawley et al., 2006) were previously identified in *S.* Typhimurium strain SL1344 background. Additionally, *Salmonella* genes required for gastrointestinal colonization of pig, calf and chicken were identified in *S.* Typhimurium strain ST4/74 (Chaudhuri et al., 2013), and those for intraperitoneal infection of mice (Sp-Liv) were reported in *S*. Typhimurium strain 14028s background (Silva-Valenzuela et al., 2015). The CEGs of different strain were searched for the corresponding orthologous genes in *S*. Typhimurium strain 14028s background using Prokaryotic Genome Analysis Tool (PGAT) (Brittnacher et al., 2011). To get insight into the phenotypic basis of CEGs required for *in vivo* intestinal colonization of pig, calf and chicken, these CEGs were compared with CEGs of *in vitro* host stressors found in the gut (PA, NaCl, PH3, Bile, and LB42). Similarly, for the phenotypic basis of CEGs for *in vivo* systemic infection (A-Mice, MΦ, P-Mice and Sp-Liv) were compared to *in vitro* macrophage stressors (H_2_O_2_, NaCl, Starvation, dLB, and PH3). Only the CEGs that were common between at least one of the *in vitro* host stressors and at least one of *in vivo* infection were identified and included for the comparative analysis.

## Results and Discussion

### Overall evaluation of resulting Tn-seq profiles

We have constructed a highly saturated transposon mutant library of *S*. Typhimurium 14028s with approximately 350,000 transposon mutants created via transformation of EZ-Tn5 transposome complex to electrocompetent cells. The complex Tn5 library, input pool 1 (IP1) was then subjected to negative selection under the *in vitro* stress conditions encountered during enteric and systemic infection as described in Materials and Methods. Input pool 2 (IP2) was the technical replicate of IP1 to evaluate the reproducibility of our Tn-seq method (Figure 1). Tn-seq amplicon library for Illumina sequencing was prepared for each of the input and output pools (Figure S1A and S1B). This efficient Tn-seq protocol was developed in our laboratory that offers distinctive advantages over other Tn-seq library preparation methods, including a low amount (~100 ng) of DNA required, and no need for physical shearing or restriction digestion(Dawoud et al., 2014; Karash et al., 2017; Kwon et al., 2016; Mandal and Kwon, 2017).

Illumina sequencing using HiSeq 3000 produced 163,943,475 reads from a single flow cell lane. The raw reads were demultiplexed allowing a perfect match for the barcodes used (Table S1) with exception of up to two mismatches within Tn5 mosaic end (ME) using a custom Perl script. H_2_O_2_ (19,250,956) had the highest number of reads followed by IP1 (10,842,764), Starvation (9,518,226), IP2 (6,345,173), LB (5,004,934), PH3 (3,841,401), PA (2,113,033) and NaCl (1,970,072) (Figure 2A).

**Figure 2.**
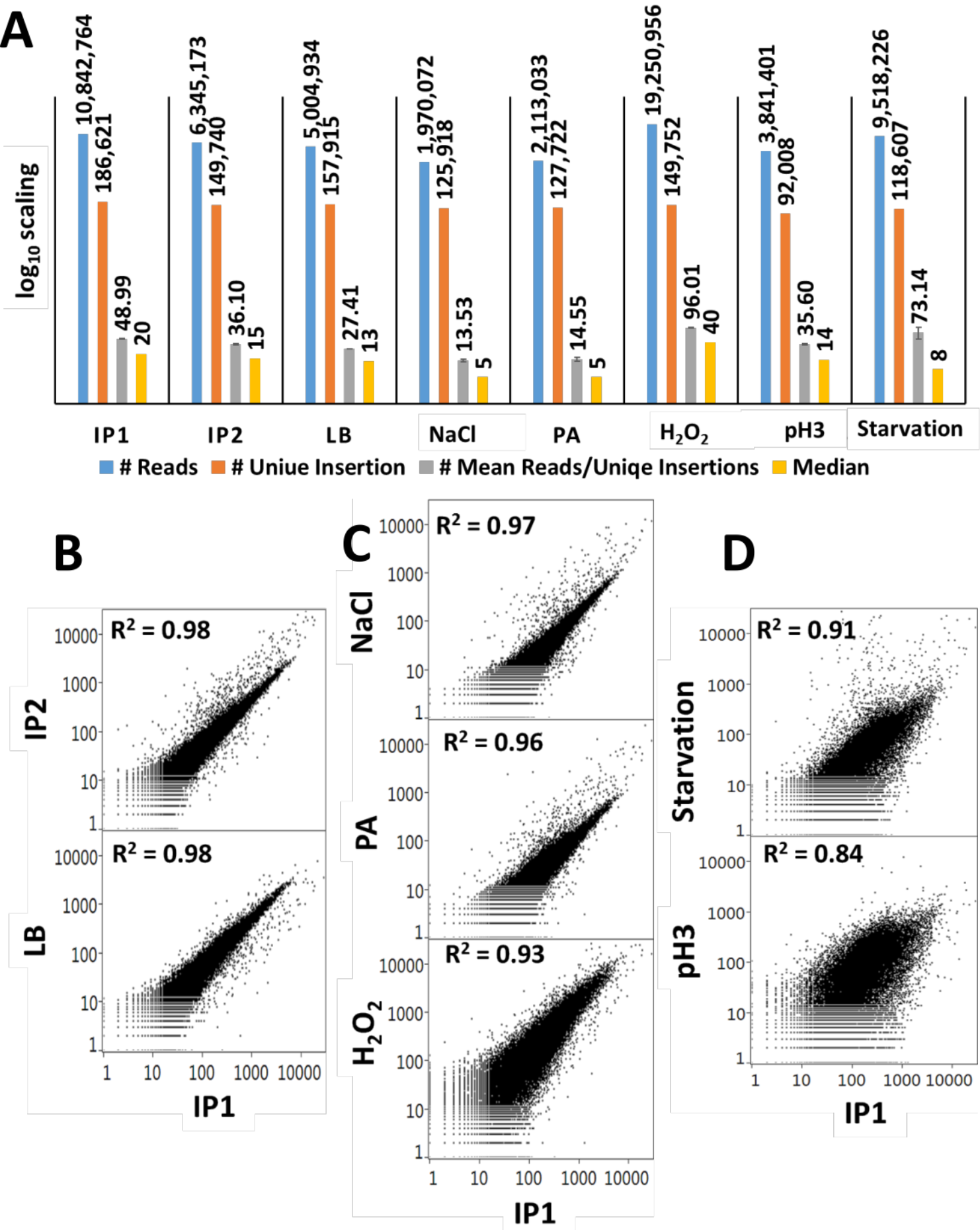
Summary of Illumina sequencing reads and correlation between Tn5 mutant libraries. A) Bar graph shows the number of Illumina sequencing reads distribution in each Tn5 libraries after sorting according to the barcode (blue color), unique insertions (orange color), mean reads per unique insertions (grey color), and median reads for each unique insertions (yellow color). B) Scatter plot displays the Spearman correlation (R^2^) among the Tn5 mutant libraries based on read count per 100 bp window across the genome (p < 0.0001).

After demultiplexing, Illumina reads were trimmed of barcode and transposon sequences. The Tn5-junction sequences of 20bp were extracted and mapped to the complete genome of *S.* Typhimurium 14028s (NC_016856.1) using Bowtie. The overall alignment rate throughout all Tn5 libraries were 85.19% (SE ± 1.79). Additionally, we looked for the unique insertion sites in the genome in each library. IP1 had the highest number of unique insertions (186,621) followed by LB (157,915), H_2_O_2_ (149,752), IP2 (149,740), PA (127,722), NaCl (125,918), Starvation (118,607) and PH3 (92,008) (Figure 2A). Similarly, H_2_O_2_ had the highest average read per unique insertion site in the genome (96.007 ± 1.11) with 40 median reads, whereas NaCl had the lowest (13.53 ± 0.99) with 5 median reads (Figure 2A).

Pre-aligned reads of the Tn5 library in default SAM mapping file format were fed to ‘Analysis of high-Resolution Transposon-Insertion Sequences Technique’ (ARTIST) pipeline (Pritchard et al., 2014). Tn5 insertions were mapped into 100 bp genome-wide windows. We observed the highest Spearman correlation coefficients (a commonly used numerical measure to describe a statistical relationship p between two variables) between IP1 and IP2, and IP1 and LB (0.98, p < 0.0001). However, there was lower Spearman correlation of IP1 with NaCl (0.97, p < 0.0001), PA (0.96, p < 0.0001), and H_2_O_2_ (0.93, p <0.0001). We observed the lowest correlation of IP1 with PH3 and Starvation (0.84 and 0.91 respectively, p < 0.0001) (Figure 2B). These relationships corroborate well with the Tn5 library selection strategies employed, with a higher correlation for the selections based on growth fitness (NaCl, PA, and H_2_O_2_) and a lower correlation for the selections based on survival (PH3 and Starvation).

Besides, we looked for the occurrence of any hot spots of Tn5 insertion in the sample libraries. We found an even distribution of Tn5 insertion reads across the libraries throughout the genome. Some of the genomic regions lacking insertions have white stripes that are clearly visible (Figure S2) across all the samples that represent essential loci in the *S.* Typhimurium 14028s genome.

### Identification of Conditionally essential genes (CEGs)

In this study, we used two strategies to identify conditionally essential genes (CEGs) of *S*. Typhimurium to overcome host stressors. The first strategy was a negative selection of complex Tn5 mutant libraries based on growth fitness for mild stressors (3% NaCl, 100 mM propionate, 1 mM H_2_O_2_) and the second one was based on survival of Tn5 mutant libraries for harsher stressors (12 days starvation and PH3) as shown in Figure 1.

The ARTIST pipeline can identify if genes are entirely essential or domain essential in a given condition. In our study only a few of the genes were identified as domain essential and the majority of them were entirely essential. For simplicity, we assigned both categories of the genes entirely essential and domain essential into one category, conditionally essential genes (CEGs). We deliberately compared the each of the output pool PA, NaCl, and H_2_O_2_ with both IP1 and LB separately. As expected, most of the CEGs were overlapped with these two comparisons. For the conditions PA, NaCl, and H_2_O_2_, we considered the common set of identified CEGs via the comparison of output library with both IP1 and LB as CEGs for each condition. However, the output libraries for PH3, and Starvation were compared only with IP1 because the selection of the Tn5 library was based on survived mutants and the mutant cells did not multiply during selection in liquid media.

We identified an overlapping set of 339 CEGs that are required for fitness of *S.* Typhimurium 14028s in at least one of the five conditions (Figure 3A). Starvation had the highest CEGs (241), followed by PH3 (103), NaCl (60), H_2_O_2_ (40) and PA (19) as shown in Table S2 and S3. This might likely reflect that starvation is a severe stressor involving diverse genetic pathways for survival, while PA is a mild stressor for the fitness of *S.* Typhimurium. More than a half of CEGs were on the lagging strand (56.63%), which is somewhat contrary to the responsive genes in *Escherichia coli* and *Streptococcus pneumoniae* (Nichols et al., 2011; van Opijnen and Camilli, 2012). We assigned a functional role to 96 CEGs that were putative proteins and 21 CEGs belonging to hypothetical proteins. The stress tolerant proteins commonly identified in at least 2 of the *in vitro* stressors included ATP synthase, a transcriptional regulator, 3-dehydoroquinate synthase, site-specific tyrosine recombinase *xerC*, flavin mononucleotide phosphatase, ribulose-phosphate 3-epimerase, and DNA-dependent helicase II among others (Table S2 and S3).

**Figure 3.**
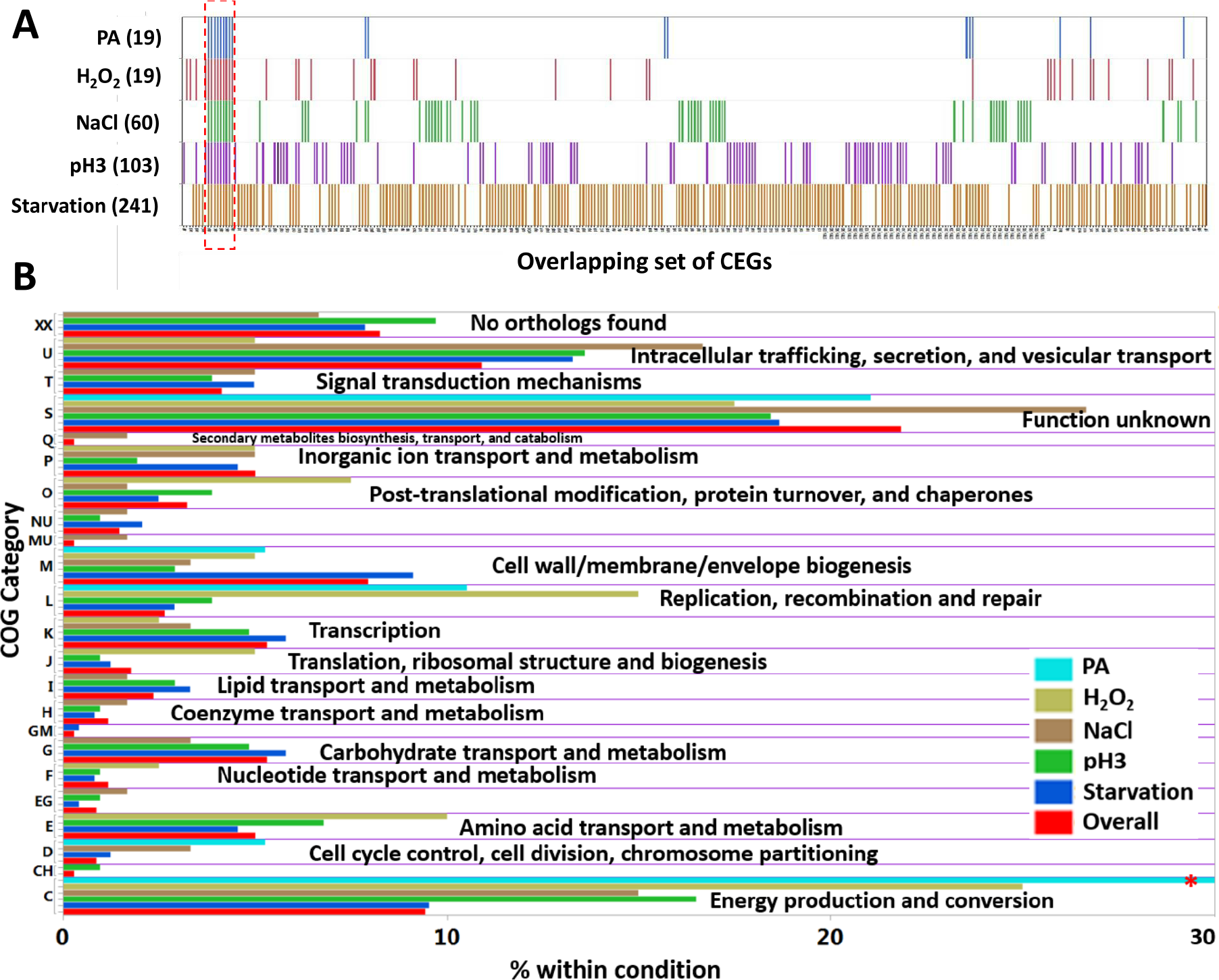
Conditionally essential genes (CEGs) of *S*. Typhimurium 14028s and cluster of orthologous groups (COG) A) Distribution of the overlapping set of 339 CEGs identified in the 5 conditions. Numbers inside the bracket indicate number of CEGs identified. Red dashed box indicates the CEGs (ATP synthase genes) common to all 5 conditions. B) Functional assignments of CEGs into COG category. Overall is the COG assigned to all the 339 CEGs. (Red asterisk (*): Abundance of COG C in PA was 57.89 %).

Intriguingly, we found many genes in the *Salmonella* pathogenicity islands (SPI) were required for fitness in the presence of the *in vitro* stressors used in this study. Numerous genes in SPI-1, SPI-2, SPI-3, SPI-5, SPI-6, and SPI-11 were required for resistance against Starvation (n=68), NaCl (n=28), and PH3 (n=27) (Table S4). However, no SPI genes were required for fitness in PA and H_2_O_2_. SPI-5 and SPI-11 genes were only conditionally essential in PH3 (n=4 and 6, respectively), while SPI-3 genes in NaCl (n=7) and SPI-6 genes in starvation (n=7). Tn-seq profiles for SPI-1 region is shown in Figure S3A as an example.

For a broader insight into pathways involved in stress resistance, we assigned each CEGs to the cluster of orthologous groups (COG) using eggNOG database (evolutionary genealogy of genes: Non-supervised Orthologous Groups) (Jensen et al., 2008). The CEGs having top hit for the COG in the *S*. Typhimurium LT2 were kept and CEGs with no orthologous group were allotted to group XX (Figure 3B; Table S3). In overall, 21.83% of CEGs belonged to category “function unknown” followed by “intracellular trafficking, secretion, and vesicular transport” (10.91%), “energy production and conversion” (9.44%), and “no orthologs found” (8.26%) among others. A substantial portion of CEGs (30.6%) falling into either “function unknown or “no orthologs found” shows that our data set is rich in novel genotype-phenotype relationships.

Additionally, we were interested to see if any CEGs identified in our study fell into the essential genomes of *S*. Typhimurium in other strain backgrounds. Essential genomes of *S.* Typhimurium strain SL3261 (selected on LB agar) (Barquist et al., 2013) and *S.* Typhimurium strain LT2 (selected on rich medium) (Knuth et al., 2004; Zhang et al., 2004) were compared with the CEGs of *S*. Typhimurium 14028s identified in this study. Genes in different strain background were looked for the corresponding orthologous genes in *S*. Typhimurium 14028s background. Interestingly, 10 and 15 CEGs in this study were shared with the essential genes of *S*. Typhimurium SL3261 and LT2, respectively (Table S5; Figure S4). This indicates that these genes that are essential in other strain backgrounds are dispensable in *S.* Typhimurium 14028s strain background.

### Molecular and phenotypic basis of CEGs in *S*. Typhimurium

Next, we delved into the genetic and biochemical mechanisms related to the CEGs identified in our study. For convenience, we split the section into specific CEGs, required for fitness in only one stressor, and common CEGs shared in at least two stressors out of five host stressors.

#### CEGs specifically required for propionate (100 mM PA) stress resistance

CEGs specific for fitness of *S.* Typhimurium in propionate were *yiiD* and *sdhAD*. YiiD is a putative acetyltransferase protein (Read coverage shown in Figure 4C). Acetylation, a post-translation modification of protein was previously shown to enable prokaryotes to increase stress resistance (Ma and Wood, 2011). Additionally, succinate dehydrogenase flavoprotein (*sdhA*) and cytochrome b566 (*sdhD*) subunit proteins were up-regulated by intestinal SCFA in *S.* Typhimurium (Lawhon, 2002). Chowdhury and Shimizu (2008) reported that *sdhA* in the tricarboxylic acid cycle (TCA) were highly induced during temperature upshift in *E. coli* (Hasan and Shimizu, 2008).

**Figure 4.**
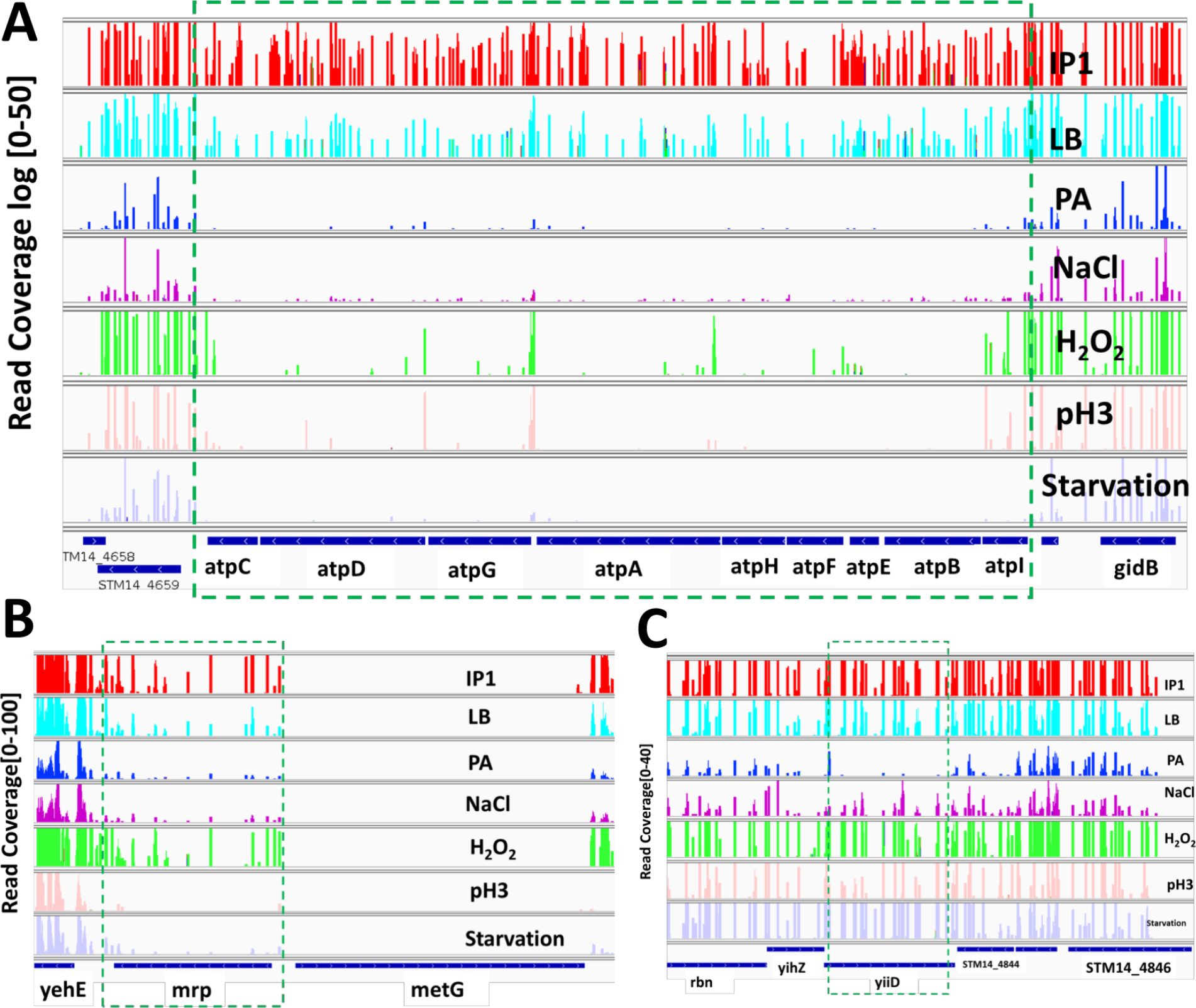
Tn-seq profiles for selected genes across 7 conditions. Y-axis: Numbers in the bracket indicates the raw read coverage. A) ATP synthase genes conditionally essential in all the 5 conditions (PA, NaCl, H_2_O_2_, PH3 and Starvation. B) Gene *mrp* essential in PH3 and Starvation. C) Gene *yiiD* essential in PA only.

#### CEGs specifically required for osmotic (3% NaCl) stress resistance

Twenty-six resistance genes of *S.* Typhimurium were required for fitness in osmotic stress (3% NaCl) alone. Protein-protein network analysis using STRING database (http://string-db.org) against *S. enterica* LT2 showed three distinct clustering of genes, SPI-3 (*mgtBC*, *misL*, *cigR*, *slsA*, *fidL* and *marT*), two-component system (*dcuBRS*) and sodium ion transport (*yihPO*) along with other nodes (http://bit.ly/2bCKGVG). SPI-3 genes are important for intracellular replication inside phagosome where *Salmonella* experience hyperosmotic stress (Schmidt and Hensel, 2004). The virulence proteins *mgtC* and *mgtB*, Mg^2+^ transporter were expressed five-fold when *S.* Typhimurium was exposed to 0.3 M NaCl (Lee and Groisman, 2012). MisL, an autotransporter protein is an intestinal colonization factor (activated by *marT*, a transcriptional regulator) that binds to extracellular matrix fibronectin in an animal host and is also involved in adhesion to plant tissue (Dorsey et al., 2005; Kroupitski et al., 2013). Deletion of *cigR* in *S.* Pullorum resulted in a significantly decreased biofilm formation and increased virulence (Yin et al., 2016). Additionally, Figureueira et al., showed Δ*cigR* strain of *S*. Typhimurium had attenuated replication in mouse bone marrow-derived macrophage (Figueira et al., 2013).

*yihPO* genes are essential for capsule assembly that is required by *Salmonella* for environmental stress persistence such as desiccation (Gibson et al., 2006). The absence of *ompL* (ortholog of *yshA*) leads to solvent hypersensitivity as it helps in the stabilization of cell wall integrity protecting from solvent penetrance as a physical barrier (Murinova and Dercova, 2014). In *E. coli*, the genes under the control of *dcuS*-*dcuR*, a two-component system, were not affected upon a hyperosmotic shock (Weber and Jung, 2002). However, *dcuBRS* were conditionally essential in *S*. Typhimurium for fitness during osmotic stress. Putative cytoplasmic protein (STM14_4542, STM14_4828, and STM14_5175), putative inner membrane protein (STM14_4824 and STM14_5184) and putative hydrolase (STM14_4823) were also required for osmotic stress tolerance.

#### CEGs specifically required for oxidative (1 mM H_2_O_2_) stress resistance

We identified 16 specific resistance genes required for fitness of *S*. Typhimurium in the presence of 1 mM H_2_O_2_ and the functional protein association network analysis among the genes was constructed using STRING against *S. enterica* LT2 (http://bit.ly/2bsVKXF). Major resistance genes were those involved in two-component system (*glnD*, *rpoN*, *arcA* (STM4598), and *arcB* (STM3328)), DNA recombination (*recJ*, and *xerD*), and metal ion transport (*corA*, and *trkA*).

Hydrogen peroxide kills *E. coli* cells with two distinct modes, mode-1 killing occurs at a lower concentration of H_2_O_2_ due to DNA damage and mode-2 killing occurs at a higher concentration of H_2_O_2_ due to damage of other structures like proteins and lipids(Imlay and Linn, 1986).

Nucleic acid metabolic process genes involved in oxidative stress resistance were *recJ*, *xerD*, *sun*, and *rpoN. RecJ* protein, a single-stranded DNA (ssDNA)-specific 5’-3’ exonuclease/deoxribophophodiesterase, plays a role in homologous recombination, mismatch repair, and base excision repair (Wakamatsu et al., 2011). In *E. coli*, *xerD* knockout mutants are hypersensitive to tightly bound DNA-protein complexes (TBCs) that block replication forks *in vivo* (Henderson and Kreuzer, 2015). *RpoN*, the alternative sigma factor 54 (σ^54^), an important regulator of stress resistance and virulence genes in many bacterial species (Riordan et al., 2010). σ^54^ is involved in carbon/nitrogen limitation, nucleic acid damage, cell envelope, and nitric oxide stress (Hartman et al., 2016). However, Hwang *et al.*, 2011 found that *rpoN* mutant in *Campylobacter jejuni* was more resistant to 1 mM H_2_O_2_ (Hwang et al., 2011).

Besides, cellular component genes crucial for fitness in H_2_O_2_ stress were *dsbC*, *glmS*, *trkA*, *corA* including *sun* and *xerD. DsbC*, a protein essential for disulfide bond isomerization in the periplasm, has a new role in *E. coli* in protection against oxidative stress (Denoncin et al., 2014). In *E. coli GlmS* plays an important role in cell wall synthesis thus providing protection against cell envelope stress response (Zhou et al., 2009). HscB, a chaperone-encoding gene is upregulated after exposure to oxidative stress in *Burkholderia pseudomallei* (Jitprasutwit et al., 2014). YbgF, an outer membrane vesicle protein, increases the survival of bacteria during exposure to stress or from toxic unfolded proteins by releasing the unwanted periplasmic component (Gogol et al., 2011).

#### CEGs specifically required for higher acidic (pH 3) stress resistance

We found 49 specific stress resistance genes required only for survival of *S*. Typhimurium in extreme acidic condition (pH 3) among other stressors. Formate dehydrogenase (*fdoHI*, and *fdhDE*) curli proteins (*csgBDEFG*), virulence and envelope proteins (SPI-2: *orf245*, *orf408*, *ssaB*; SPI-5: *pipBC*, *sopB*, and SPI-11: *envEF*, *pagCD*, *msgA*, STM14_1486 where *ssaB*, *pipB*, and *sopB* are effector proteins), and biopolymer transport protein (*exbD* and *exbB*) were clustered in functional protein association network analysis using STRING (http://bit.ly/2bCLVnL).

Formate dehydrogenase catalyzes the oxidation of formate (HCOO-) to CO_2_ and H^+^. The released electrons from this reaction are used by two cytoplasmic protons to form dihydrogen thus consuming net protons, consequently, counteracting acidification (Leonhartsberger et al., 2002). Curli are major complex extra-cellular proteinaceous matrix produced by *Enterobacteriaceae* that helps pathogenic bacteria like *Salmonella* in adhesion to surfaces, cell aggregation, and biofilm formation (Barnhart and Chapman, 2006). Acidic pH strongly enhances biofilm formation in *Streptococcus agalactiae* (D’Urzo et al., 2014). We hypothesize that curli fibers might potentially protect bacteria from severe acid stress through the physical barrier and likely by the generation of alkaline compounds as in oral biofilms (Cotter and Hill, 2003). PhoP regulates SPI-11 genes such as *envEF*, *pagCD*, and *msgA* where later three are required by *Salmonella* to survive low pH within macrophage (Gunn et al., 1995; Lee et al., 2013). In *Helicobacter pylori*, only the organism to colonize in the acidic human stomach, *ExbB*/*ExbD*/*TonB* complex is required for acid survival and periplasmic buffering (Marcus et al., 2013). Additionally, survival of *ΔexbD* was diminished compared to wild type at pH 3 in *E. coli* (Ahmer et al., 1995). The *metC* gene encoding a key enzyme in methionine biosynthesis, required for the generation of homocysteine, pyruvate, and ammonia, play a crucial role in bacterial acid stress responses (Reid et al., 2008).

#### CEGs specifically required for starvation stress resistance

Out of 261 *Salmonella* fitness genes essential for starvation stress, 160 genes were explicitly important for resistance against starvation stress among the five infection-relevant conditions in this study (http://mcaf.ee/k0uhrm). Major enriched gene pathways were oxidative phosphorylation, pathogenesis, two-component system, and lipopolysaccharide biosynthetic process among others. NADH dehydrogenase, the first component of the respiratory chain, subunit proteins (*nuoCEFGHLMN*) were required for fitness of *Salmonella* during long-term carbon starvation. *Salmonella* defective in NADH dehydrogenase enzyme exhibits defective energy-dependent proteolysis during carbon starvation (Archer et al., 1993). Proteolysis of unbound or unemployed proteins helps bacteria to access nutrients as an important survival strategy during carbon starvation (Michalik et al., 2009). SPI-1 (*hilACD*, *iagB*, *invH*, *orgAC*, *prgHIJK*, STM14_3500, and STM14_3501) and SPI-2 (*ssaMNOPQRSTV*, *sscB*, and *sseDEF*) encoding type III secretion system (T3SS) and SPI-6 (*safABCD*, *sinR*, STM14_0359, and *ybeJ*) encoding type VI (T6SS) secretion system were required for *in vitro* survival in long-term starvation stress. *Salmonella* usually requires SPI-1 genes for the invasion of intestinal epithelial cells (Klein et al., 2000). HilACD regulates SPI-1 invasion gene expression during multiple environmental conditions including stationary phase, pH, osmolality, oxygen tension, and short chain fatty acids (Olekhnovich and Kadner, 2007). SPI-2 genes are expressed under *in vitro* starvation conditions indicating the use of nutritional deprivation as a signal (Hensel, 2000). T6SS has been hypothesized to confer a growth advantage to bacteria in environmental niches where bacterial competition for nutrient is critical for survival (Brunet et al., 2015).

Two-component systems (TCs), a basic stimulus-response coupling mechanism, enable microbes to respond to various stimuli such as pH, osmolality, quorum signals, or nutrient availability and regulate their cellular functions (Freeman et al., 2013). TCs required for fitness during starvation conditions were *envZ*/*OmpR*, *cpxA*/*cpxR*, sensory histidine kinase protein (*phoQ*), and kdpD (Figure S3B). EnvZ/*OmpR* regulates the synthesis of porin proteins (*ompF* and *OmpC*) that are important for the survival of *E. coli* in sea water under starvation stress condition (Darcan et al., 2009). It is believed that carbon starvation causes cell envelope stress. Bacchelor et al., (2005) found *cpxA*/*cpxR* in *E. coli* regulates the expression of prions *ompF* and *ompC*, a major component of the outer membrane. However, Kenyon et al., (2002) showed the starvation stress of *S.* Typhimurium do not require cpxR-regulated extra-cytoplasmic functions (Batchelor et al., 2005; Kenyon et al., 2002). PhoQ and *kdpD* plays a role in Mg^2+^ and K^+^ homeostasis respectively, critical to the virulence and intracellular survival of *S*. Typhimurium (Freeman et al., 2013; Kato and Groisman, 2008).

The outer membrane of Gram-negative bacteria contains phospholipids and lipopolysaccharides (LPS). LPS molecules act as a permeability barrier to prevent the entry of toxic compounds and allow the entry of nutrient molecules (Schakermann et al., 2013). LPS biosynthetic process genes required for fitness in starvation conditions were *rfbABCD*, *rfbUNMKP, galF*, *udg*, *wzxE*, and *wzzB*. Starvation of carbon energy source activates envelope stress response in *S.* Typhimurium (Rowley et al., 2006). Additionally, *pstSCAB* coding for the Pst ABC transporter catalyzes the uptake of inorganic phosphate (Lüttmann et al., 2012). Mutations in the Pst system results in structural modifications of lipid A and an imbalance in unsaturated fatty acids consequently leading to increase in outer membrane permeability making *E. coli* more vulnerable to environmental stresses including antimicrobial peptide and low pH (Lüttmann et al., 2012).

Additional genes required for starvation stress resistance were *aroGH*, ytfMNP (ytfM - outer membrane protein), *stcB* (putative periplasmic outer chaperone protein). Furthermore, other envelope proteins were outer membrane lipoproteins (*stcD* and *yifL*), putative outer membrane proteins (*stcC*, STM14_0404, and ytfM), and putative inner membrane proteins (STM14_0398, STM14_0402, STM14_2763, STM14_4741, STM14_4742, STM14_4745, STM14_4880, *ydiK* and *yjeT*). Similarly, putative cytoplasmic proteins required for starvation stress were STM14_2759, STM14_4743, STM14_5374, *ydiL*, and *ytfP*.

#### CEGs required for tolerance to multiple stressors

We found 12 *Salmonella* genes required for stress resistance in either three or four of the *in vitro* host stresses in our study as shown in STRING protein-protein interaction network (http://bit.ly/2btx1zg). The enriched GO biological process / KEGG pathways were ncRNA processing (*gidAB* and *mnmE*), DNA metabolic process (*dam*, *uvrD* (SOS response), *xerC*), and biosynthesis of amino acids (*aroB* and *rpe* - microbial metabolism in diverse environments). In addition, other responsive proteins include ATP synthase subunit protein (*atpI*), putative permease (STM14_4659), inner membrane protein (*damX*), and flavin mononucleotide phosphatase.

*DamX*, *dam*, *rpe*, *aroB*, *uvrD*, and *yigB* were required for fitness in PH3, Starvation, and H_2_O_2_. Disruption of *damX* in *S. enterica* causes bile sensitivity (López-Garrido and Casadesús, 2010). DNA adenine methylation gene (*dam*) plays an important role in bacterial gene expression and virulence (Low et al., 2001). Dam mutants of *S. enterica* are extremely attenuated in mouse (Jakomin et al., 2008). The gene *aroB* encodes dehyroquinate synthase, a part of shikimate pathway, is essential for bacteria and absent in mammals (de Mendonca et al., 2007). In prokaryote species, *uvrD* is involved in maintaining genomic stability and helps DNA lesion repair, mismatch repair, nucleotide excision repair and recombinational repair (Kang and Blaser, 2006). Overproduction of *yigB* produced higher-level persister, cells that exhibit multidrug tolerance, in *E. coli* (Hansen et al., 2008). However, deletion of *gidB* (glucose-inhibited division gene B) confers high-level antimicrobial resistance in *Salmonella* and has compromised overall bacterial fitness compared to wildtype (Mikheil et al., 2012). GidA (together with *mnmE*) is responsible for the proper biosynthesis of 5-methylaminomethtyl-2-thouridine of tRNAs and deletion causes attenuation in bacterial pathogenesis (Shippy and Fadl, 2014b).

### ATP synthase genes are obligatory for *Salmonella* fitness during *in vitro* host stressors

ATP synthase (F_1_F_0_-ATPase) is a ubiquitous enzyme largely conserved across all domains of life. All the eight genes encoding ATP synthase subunit proteins were required for fitness of *S.* Typhimurium in every 5 *in vitro* conditions of our study (Figure 3A and 4A). F_1_F_0_-ATP synthase complex is required for ATP production from ADP and Pi. ATP synthase also regulates pH homeostasis in bacteria (*Listeria monocytogenes* and *S.* Typhimurium) at the expense of ATP (Balemans et al., 2012). In *Streptococcal faecalis*, upregulation of F_1_F_0_-ATPase promotes ATP-dependent H+ extrusion under acidic conditions. However, in *E. coli* the expression of ATP synthase is decreased under acidic condition (Krulwich et al., 2011). ATP synthase in *Mycobacterium* and *Staphylococcus* has been validated as a promising target for new antimicrobial drugs (Balemans et al., 2012; Lu et al., 2014).

### Mechanistic basis of *Salmonella in vivo* fitness genes required for enteric and systemic infection

The network diagrams shown in Figure 5 and Figure 6 show all the genes that are commonly important for fitness under at least one of the *in vitro* and *in vivo* conditions. The genes that were important only either in the *in vitro* or *in vivo* conditions were excluded in the diagram.

**Figure 5.**
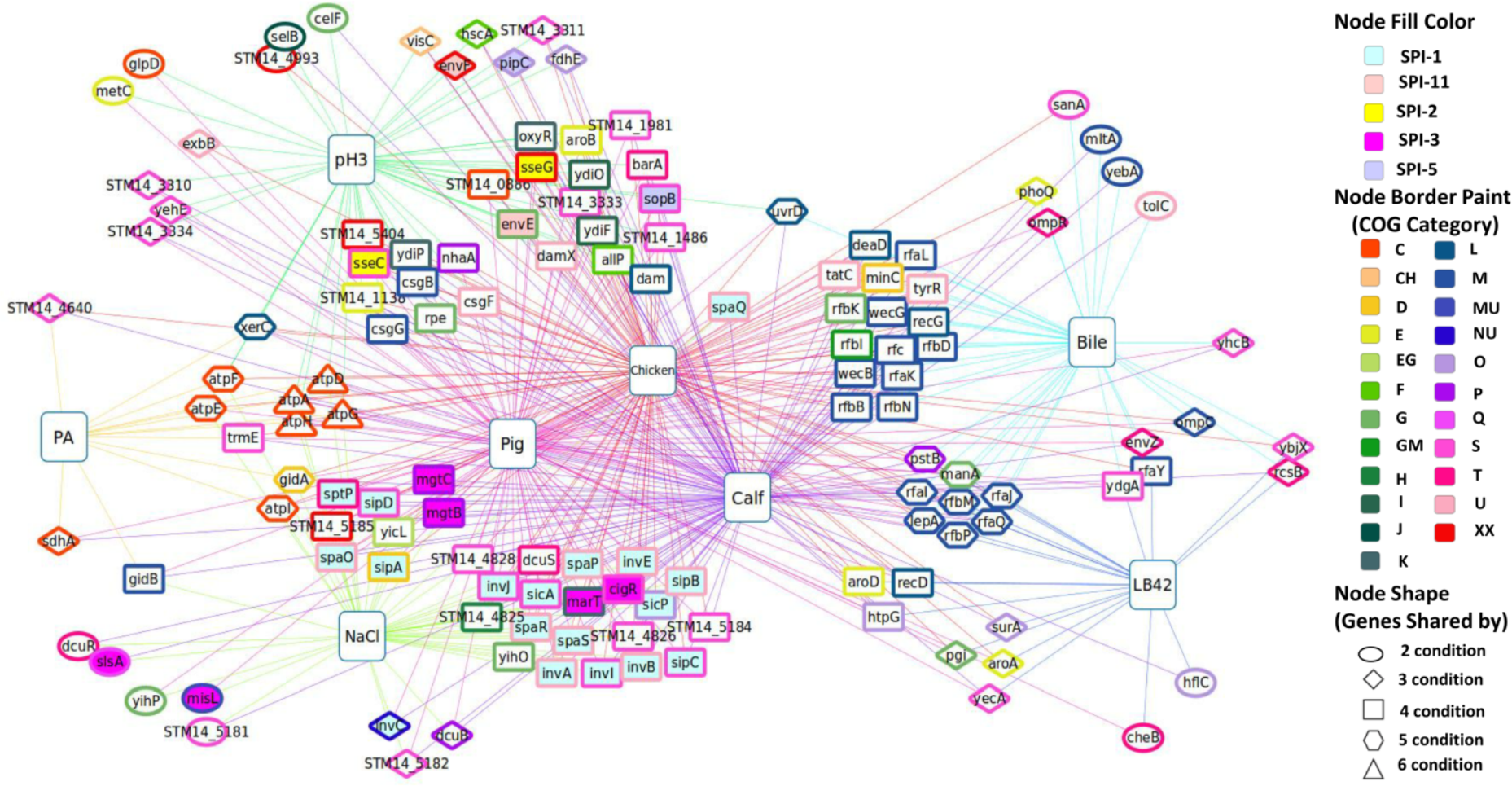
Genotype-phenotype network connections illustrating mechanistic basis of *S.* Typhimurium genetic factors required for enteric infection [*In vitro* vs *in vivo* (Enteric)] Large square nodes indicate various conditions (studies) and small nodes are fitness genes. Each node (gene) is at least shared by one of the in vitro condition i.e. stressors encountered by *Salmonella* during enteric infection (PA, PH3, NaCl, Bile, and LB42) and at least one of the *in vivo* enteric condition (Pig, Calf, and Chicken). The interactive network through the Network Data Exchange (NDEx) is available at www.ndexbio.org/#/network/027b067d-e209-11e8-aaa6-0ac135e8bacf (Pratt et al. 2015).

**Figure 6.**
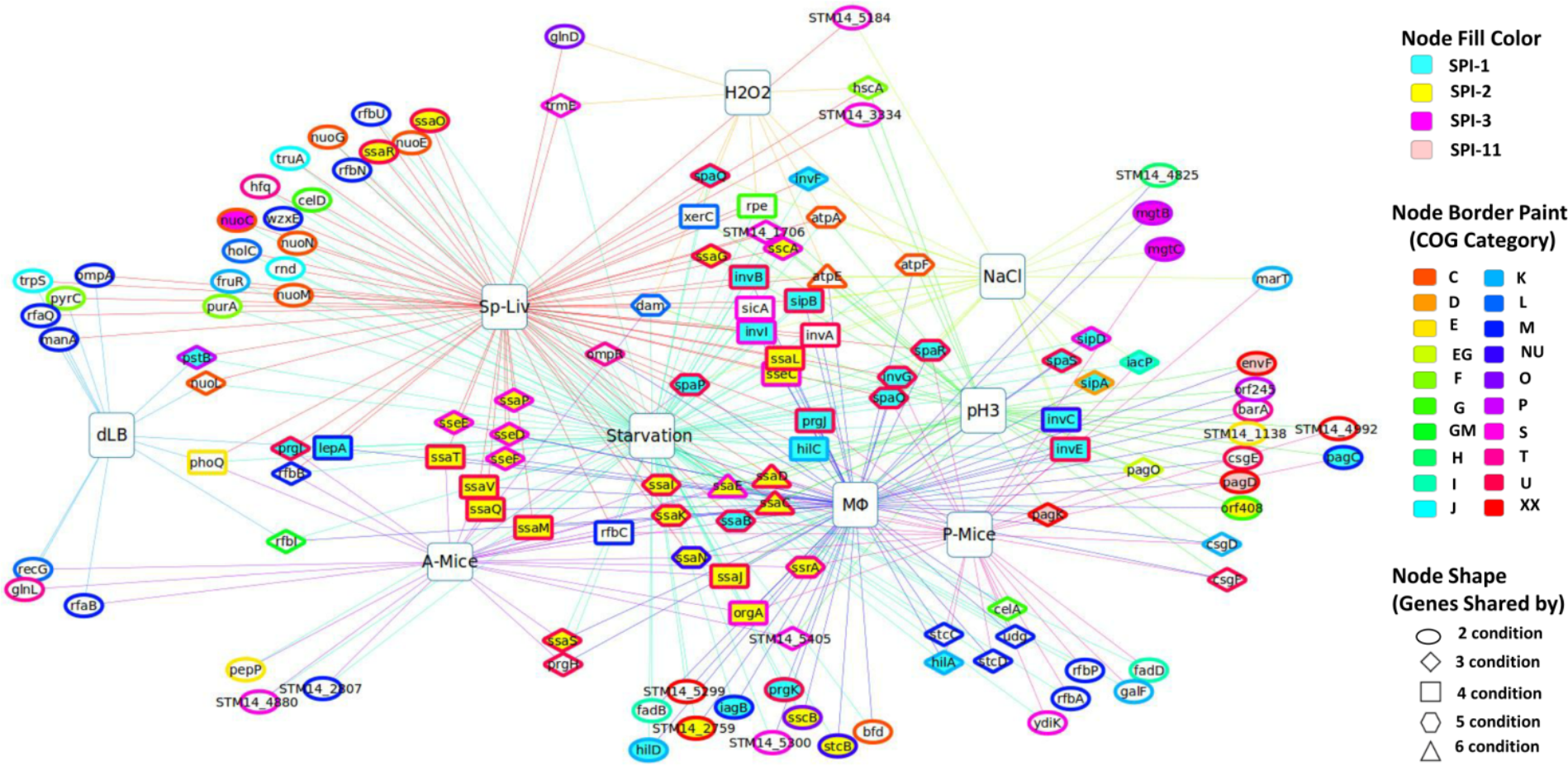
Genotype-phenotype network connections illustrating mechanistic basis of *S.* Typhimurium genetic factors required for systemic infection [*In vitro* vs *In vivo* (Systemic)] Large square nodes indicate various conditions (studies) and small nodes are fitness genes. Each node (gene) is at least shared by one of the *in vitro* condition i.e. stressors encountered by *Salmonella* inside macrophage (NaCl, H_2_O_2_, PH3, Starvation, and dLB) and at least one of the *in vivo* systemic condition (MΦ, Sp-Liv, P-Mice, and A-Mice). The interactive network through the Network Data Exchange (NDEx) is available at www.ndexbio.org/#/network/5e78ad70-e209-11e8-aaa6-0ac135e8bacf (Pratt et al. 2015).

Numerous *in vivo* fitness genes have been identified in previous studies, indicating that they are required by *S.* Typhimurium to overcome host defenses. However, for a large portion of them the mechanistic bases why they are required in particular *in vivo* niches remain unknown. The information on the common requirements of the genes shown in these networks (Figure 5 and 6) for both at least one well-defined *in vitro* stress and *in vivo* infection model is valuable in the sense that it provides novel insights on the type of selective pressures *S.* Typhimurium might be facing during infection in the host.

#### Enteric infection

We have identified an overlapping set of 135 CEGs that are commonly required to cause enteric infection in at least one of the host [pig, calf, and chicken (Chaudhuri et al., 2013)] and for fitness in one of the *in vitro* host stressors [LB42, Bile (Khatiwara et al., 2012), PH3, PA, and NaCl] encountered during enteric infection (Figure 5; Table S6). Genes in SPI-1 (*invABCEIJ*, *sicAP*, *sipABCD*, *spaOPQRS*, *sptP*) and SPI-3 (*cigR*, *marT*, *mgtBC*, *misL*, *slsA*) were required for fitness in NaCl and all host. However, genes encoding SPI-2 (*sseCG*), SPI-5(*slsA*, *pipC*) and SPI-11(*envEF*) were essential for fitness only one *in vitro* stressor PH3 and intestinal colonization in 3 hosts. Other enriched pathways were lipopolysaccharide biosynthesis (*rfaIJKLQY* and *rfbBDKMNP*), oxidative phosphorylation (ATP synthase genes and *sdhA*), and biosynthesis of amino acids (*aroABD, rpe and metC*) including others as shown in STRIN protein-protein interaction against *S. enterica* LT2 (http://mcaf.ee/wzljud).

High osmolality, low oxygen, and late log phase induce *hilA* expression *in vitro* that in turn regulates the expression of SPI-1 genes (Lostroh and Lee, 2001). Interestingly, we identified SPI-1 genes as fitness genes required for *in vitro* NaCl stressor. Similarly, lipopolysaccharide (LPS) biosynthetic process genes were enriched in LB42, Bile and in pig, calf, and chicken for fitness during enteric infection. LPS, a critical factor in the virulence of gram-negative bacterial infection is required for intestinal colonization, resistance to killing by macrophage, swarming motility, serum resistance and bile stress (Khatiwara et al., 2012; Kong et al., 2011). CsgBA (curli subunit protein) mutant of *S*. Typhimurium was attenuated to elicit fluid accumulation in bovine ligated ileal loops (Tükel et al., 2005) and are required for fitness in PH3 including *csgF* and *csgG*. Additionally, putative proteins STM14_1138, STM14_1486, STM14_1981, STM14_3333 and STM14_4826, STM14_4828, STM14_5184, STM14_5185 (hypothetical protein) were required for fitness *in vitro* acidic and osmotic stress respectively and enteric infection in the entire three host.

#### Systemic infection

We compared the CEGs that are at least shared between the one of the *in vitro* host stressors, H_2_O_2_, NaCl, PH3, Starvation and dLB (Khatiwara et al., 2012), encountered inside MΦ and *in vivo* systemic infections (MΦ (Chan et al., 2005), A-Mice (Chan et al., 2005), P-Mice (Lawley et al., 2006), Sp-Liv (Silva-Valenzuela et al., 2015) and identified an overlapping set of 130 genes (Figure 6; Table S7) shown in protein-protein interaction network using STRING (http://mcaf.ee/p34rjn). SPI-1 genes (*hilACD*, *iacP*, *iagB*, *invABCEFGI*, *orgA*, *prgHIJK*, *sicA*, *sipABC*, *spaOPQRS*) encoding TTSS were essential for fitness in NaCl, Starvation, MΦ survival and systemic infection. Additionally, SPI-2 genes (*ssaBCDEGIJKLMNOPQRSTV*, *orf245*, *orf408*, *sscAB*, *sseCDEF*, *ssrA*, STM14_1706) encoding TTSS were required for fitness in PH3, starvation, MΦ survival and systemic infection.

Similarly, SPI-3 genes (*marBCT*) were required for fitness in NaCl, MΦ survival, and persistent infection in mice (P-Mice). SPI-11 genes (envF, pagCD) were required for fitness in PH3, MΦ survival, and P-Mice.

Other than SPI genes, the majorly enriched genes were nucleic acid metabolic process (*dam*, *trpS*, *MnmE*, *truA*, *serc*, *csgD*, *ompR* and *cra*), lipopolysaccharide biosynthetic process (*rfbABCNPU*, *rfaB*, *udg*, *galF*), oxidative phosphorylation (ATP synthase genes, NADH dehydrogenase genes), two component system (*ompR*, *barA*, *phoQ*, *glnDL*, *pagKO*) among others (Figure 6). Gene *dam* was required for fitness in H_2_O_2_, NaCl, A-Mice, and Sp-Liv. XerC and *rpe* were required for H_2_O_2_, PH3, Starvation and Sp-Liv. Interestingly, *pagK* were not identified as CEG in A-Mice, P-Mice, Sp-Liv but in PH3, Starvation, and MΦ. Putative genes either essential for one of *in vitro* or *in vivo* systemic infection were STM14_1138, STM14_4880, STM14_4992, STM14_5184, STM14_2759, STM14_2807, STM14_3334, STM14_4825, STM14_5299, and STM14_5300.

### Limitations of the study

This study has some limitations. Firstly, this study was exploratory in nature. Thus, prior knowledge of CEGs regarding stress tolerance were discussed where possible rather than performing phenotypic study of single gene knockout mutants. Secondly, Tn-seq approach is prone to false positive and false negative results. However, assessment for either false positive or false negative was not performed. Additionally, domain essential genes might have increased the chance of false positive discovery which were categorized as CEGs. Lastly, the scope of comparative study was limited to the CEGs identified in this study that were compared with the previously identified CEGs either *in vitro* or *in vivo* conditions which were mainly identified using Tn-seq approach. Nevertheless, all the CEGs identified in stress conditions had significantly lower reads compared to the control group, strongly supporting true conditional essentiality of the CEGs identified in this study. Most importantly, our goal was to provide comprehensive framework for mechanistic basis of genes required for *in vivo* fitness.

## Conclusion

A recent study by Kroger et. al. (2013) presented transcriptomes of *S*. Typhimurium in 22 distinct infection-relevant environmental conditions *in vitro*. The study found induction of *Salmonella* pathogenicity islands *in vitro* conditions such as early stationary phase, anaerobic growth, oxygen shock, nitric oxide shock as well as in pH3, NaCl, bile, and peroxide shock among others (Kröger et al., 2013). However, transcription of a gene does not necessarily indicate the need of that gene function for fitness in a given particular condition. The transcript can be a leaky expression or required for fitness in the upcoming environment in a cost effective way through predictive adaptation, phenomena where bacteria are able to anticipate and pre-emptively respond to the regular environmental fluctuations (temporally distributed stimuli) that confers a considerable fitness advantage for the survival of an organism (Mitchell et al., 2009; Tagkopoulos et al., 2008). Traditionally, it is believed that “central dogma of life” i.e. flow of information from DNA to RNA to proteins are highly concordant. However, there is a modest correlation between levels of transcripts and corresponding proteins (Foss et al., 2007; Fu et al., 2009; Ghazalpour et al., 2011). Thus, functional genomics screening such as Tn-seq is expected to reveal more direct functional aspects of the genes involved in responding to the current stresses.

In this report, we were able to map genotype to phenotype links providing the mechanistic basis of the genetic requirements for fitness for an overlapping set of 221 virulence genes for *in vivo* fitness (Figure S5). These CEGs were required for fitness in at least one of the *in vitro* host stressors (PA, NaCl, PH3, Starvation, Bile, LB42 and dLB), and enteric infection (calf, chicken and pig), or systemic infection (mice including intracellular survival inside macrophage). Forty-four common CEGs were required to cause both systemic and enteric infections (*in vivo* fitness) and *in vitro* fitness (Figure S5 and Table 1). Common SPI genes for *in vivo* and *in vitro* fitness were SPI-1 (*invABCEI*, *sicA*, *sipABD*, *spaOPQRS*), SPI-2(*sseC*), SPI-3(*marT*, *mgtCB*) and SPI-11(*envF*). *Salmonella* genes other than SPI essential for fitness under *in vitro* stresses and *in vivo* survival were *atpAEF*, *lepA*, *dam*, *pstB*, *xerC*, *manA*, *phoQ*, *rfaQ*, *rfbBIP*, *rpe*, *trmE*, *rfbIP*, *ompR*, *csgF*, *recG*, *hscA*, *barA*, and putative genes STM14_1138, STM14_3334, STM14_4825, and STM14_5184 (Table 1).

**Table 1:** *Salmonella* genes required for *in vitro* and *in vivo* (enteric and systemic) fitness.

| Category | Genes | Conditions (*In vitro*,<br>enteric, and, systemic) | COG | Protein name |
| --- | --- | --- | --- | --- |
| **SPI Genes** |  |  |  |  |
| SPI-1* | *invA* | Na, S, C, P, Ch, MΦ, SL | U | needle complex export protein |
|  | *invB* | Na, S, C, P, Ch, MΦ, SL | U | secretion chaperone |
|  | *invC* | Na, S, C, P, MΦ, PM | NU | ATP synthase SpaL |
|  | *invE* | Na, S, C, P, Ch, MΦ, SL | U | invasion protein |
|  | *invI* | Na, S, C, P, Ch, MΦ, SL | S | needle complex assembly protein |
|  | *sicA* | Na, S, C, P, Ch, MΦ, SL | S | secretion chaperone |
|  | *sipA* | Na, S, C, P, Ch, MΦ | D | secreted effector protein |
|  | *sipB* | Na, S, C, P, Ch, MΦ, SL | U | translocation machinery component |
|  | *sipD* | Na, S, C, P, Ch, MΦ | S | translocation machinery component |
|  | *spaO* | Na, S, C, P, Ch, MΦ, PM, SL | U | surface presentation of antigens protein SpaO |
|  | *spaP* | Na, S, C, P, Ch, MΦ, AM, SL | U | surface presentation of antigens protein SpaP |
|  | *spaQ* | Na, S, C, P, Ch, SL | U | needle complex export protein |
|  | *spaR* | Na, S, C, P, Ch, MΦ, PM, SL | U | needle complex export protein |
|  | *spaS* | Na, S, C, P, Ch, MΦ | U | surface presentation of antigens protein SpaS |
| SPI-2* | *sseC* | pH, S, C, P, Ch, MΦ, SL | S | translocation machinery component |
| PI-3 | *marT* | Na, C, P, Ch, PM | K | putative transcriptional regulator |
| SPI-11 | *envF* | pH, C, Ch, MΦ | XX | putative envelope lipoprotein |
| **Non-SPI genes** |  |  |  |  |
| Two-component system | *ompR\** | B, S, C, P, MΦ, SL | T | osmolality response regulator |
|  | *phoQ\** | B, S, dLB, C, Ch, AM, SL | E | sensor protein PhoQ |
|  | *barA* | pH, S, C, P, Ch, MΦ | T | hybrid sensory histidine kinase BarA |
| O antigen biosynthetic process | *rfbB* | B, S, C, P, Ch, AM, SL | M | dTDP-glucose-4,6-dehydratase |
|  | *rfbP* | B, S, L4, C, P, Ch, PM | M | undecaprenol-phosphate galactosephosphotransferase/O-antigen transferase |
|  | *rfbN* | B, S, C, P, Ch, SL | M | rhamnosyl transferase |
| ATP synthase genes* | *atpA* | PA, Na, pH, H2, S, C, P, Ch, SL | C | F0F1 ATP synthase subunit alpha |
|  | *atpE* | PA, Na, pH, H2, S, P, Ch, MΦ, SL | C | F0F1 ATP synthase subunit C |

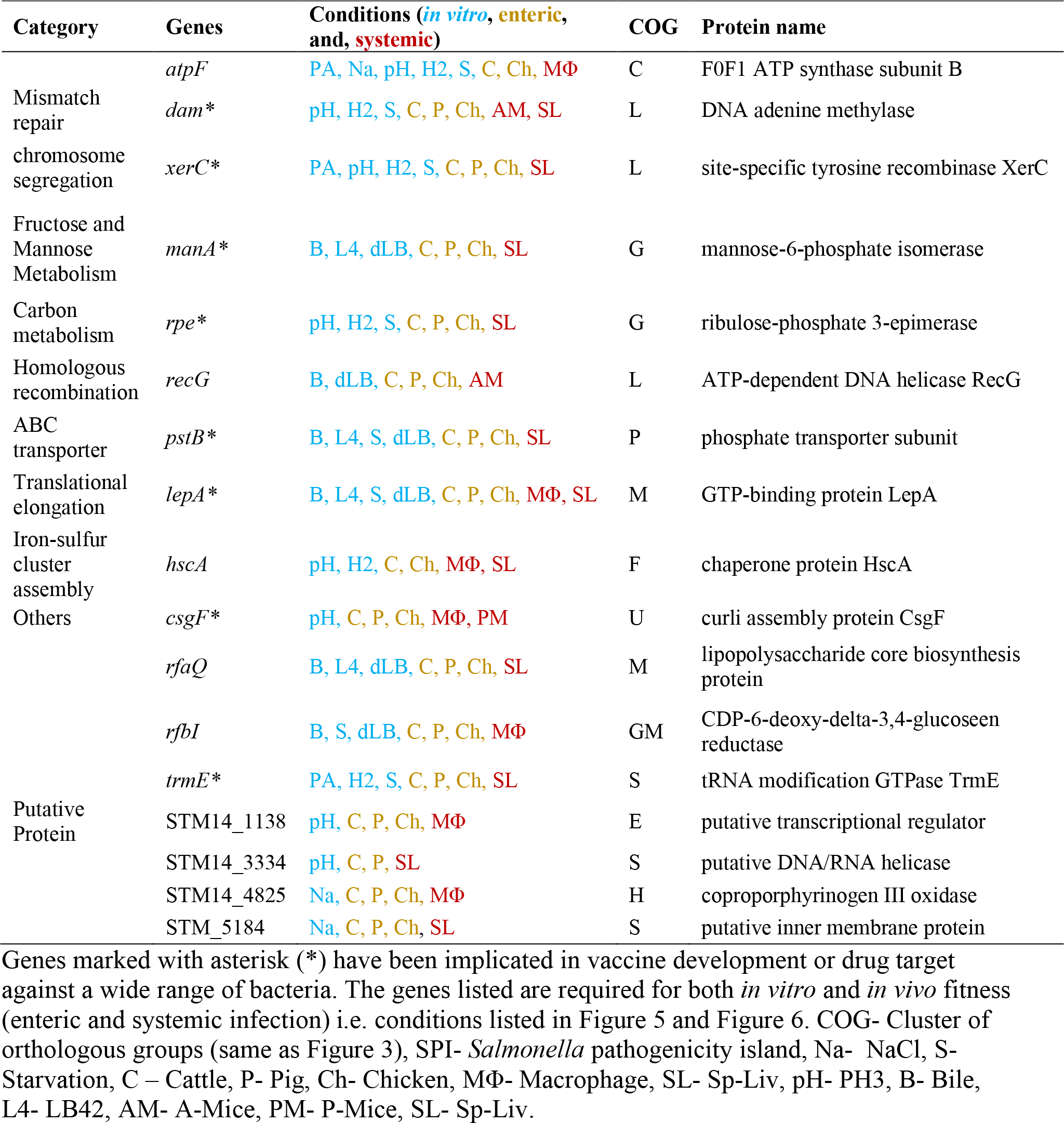

Interestingly, most of the common forty-four genes required for *in vitro* and *in vivo* (enteric and systemic infection) fitness have been implicated in vaccine or drug target development against broad spectrum of bacteria. Such as ATP synthase genes (Balemans et al., 2012; Lu et al., 2014), *dam* (Garcia-Del Portillo et al., 1999), *pstB* (Garmory and Titball, 2004), *phoQ* (Miller and Mekalanos, 1998), *ompR* (Dougan et al., 1996), *xerC* (Hur et al., 2011), and *rfbBPN* (Sturm and Timmis, 1986), *manA* (Amineni et al., 2010), *rpe* (Edwards et al., 2004), *lepA* (Patton, 2007), *csgF* (Cegelski et al., 2008), trmE (Shippy and Fadl, 2014a), and SPI-1 and SPI-2 (Matulová et al., 2012) have been used as vaccine development or drug target (Table 1). Thus, there lies a great potential to explore genes *marBCT*, *envF*, *barA*, *hscA*, *rfaQ*, *rfbI* and putative proteins STM14_1138, STM14_3334, STM14_4825, and STM_5184 as novel therapeutic and intervention strategy to curb *Salmonella* infection.

## Acknowledgements

We would like to thank Arkansas High Performance Computing Center (AHPCC) - University of Arkansas for their computational support.

## Supplemental Materials

**Figure S1.**
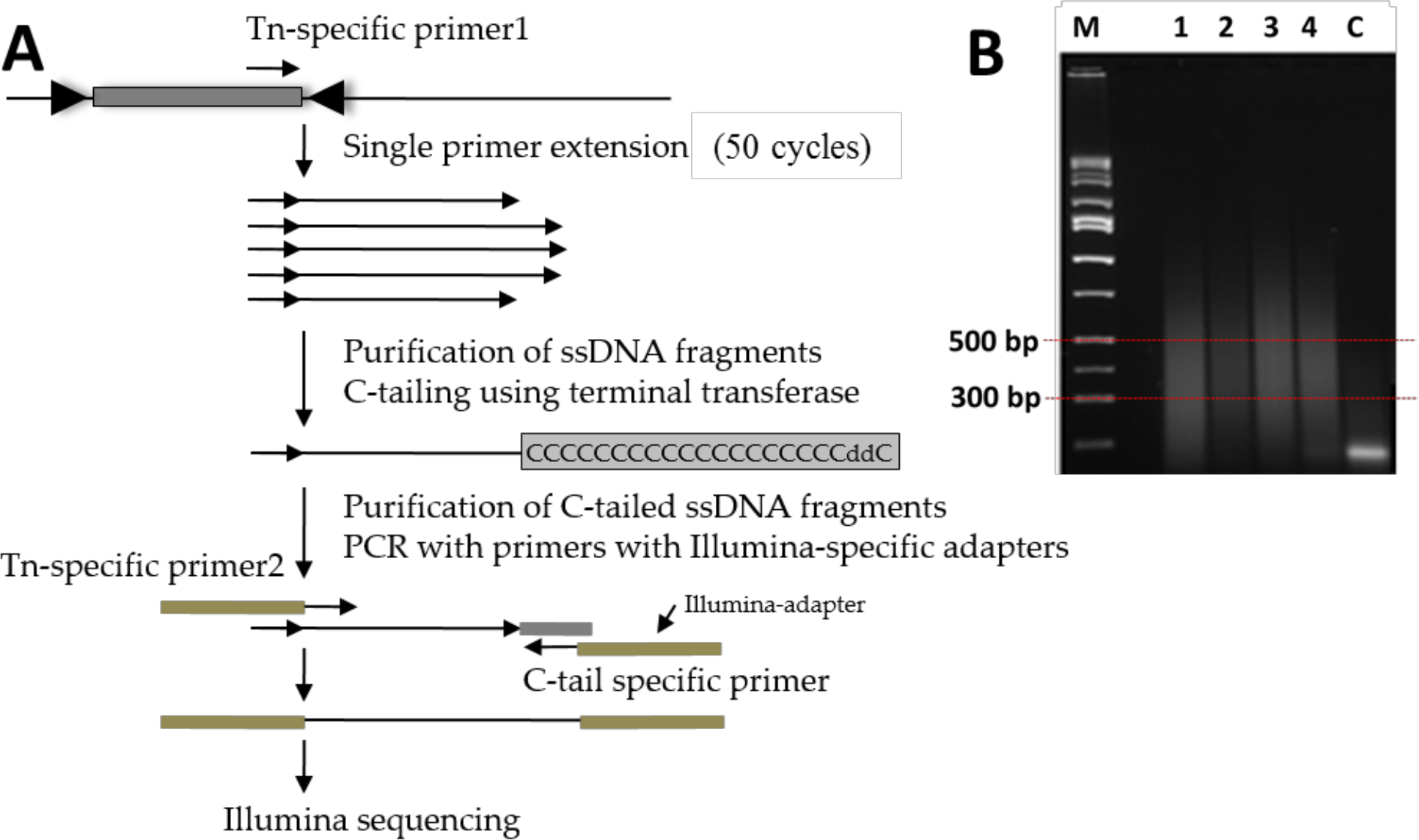
Preparation of Tn-seq amplicon library for Illumina sequencing. A) Genomic DNA of Tn5 mutant library was linealry extended using Tn-specific primer 1 (Ez-Tn5 primer3 in Table S1). Then C-tail was attached to the 3’ end of purified single-stranded DNA. The C-tailed product was purified and exponential PCR was performed using Tn-specific primer 2 (Barcoded primers in Table S1) and C-tail specific primer (HTM-Primer in Table S1) with Illumina adapter attached to primers. B) Exponentially amplified DNA was than run on 1.5% agarose gel. DNA from 300bp to 500bp was extracted from the gel and sent for Illumina sequencing. [M: Hi-Lo DNA marker; 1, 2, 3, 4: Tn5 mutant libraries; and C: negative control (gDNA of the wild type *S*. Typhimurium 14028s)].

**Figure S2.**
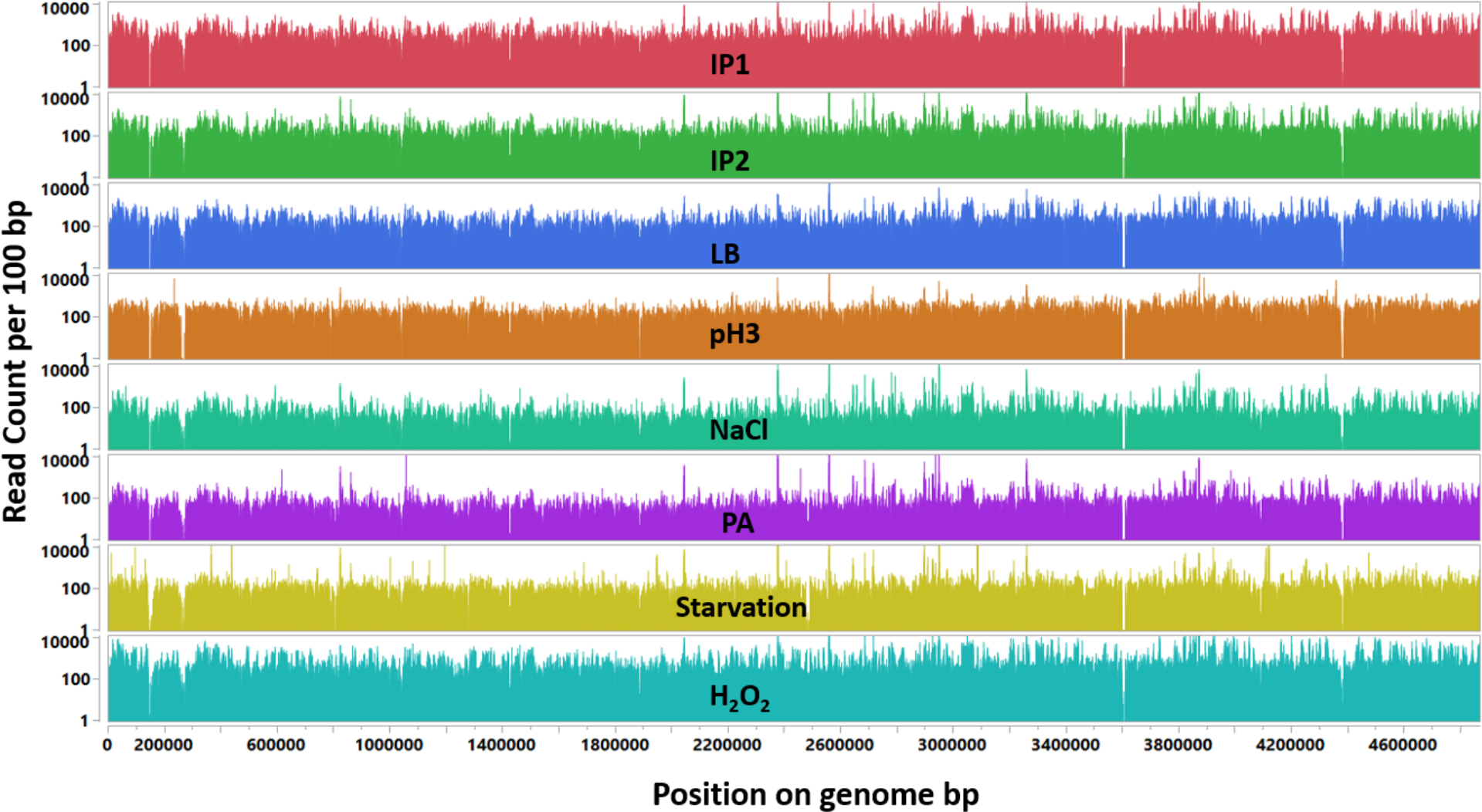
Overlay plot displays global view of genome-wide quantitative distribution of Tn5 insertion read count for all samples. X-axis: Position on the genome; and Y-axis: Number of read count per 100 bp scaled in log_10_.

**Figure S3.**
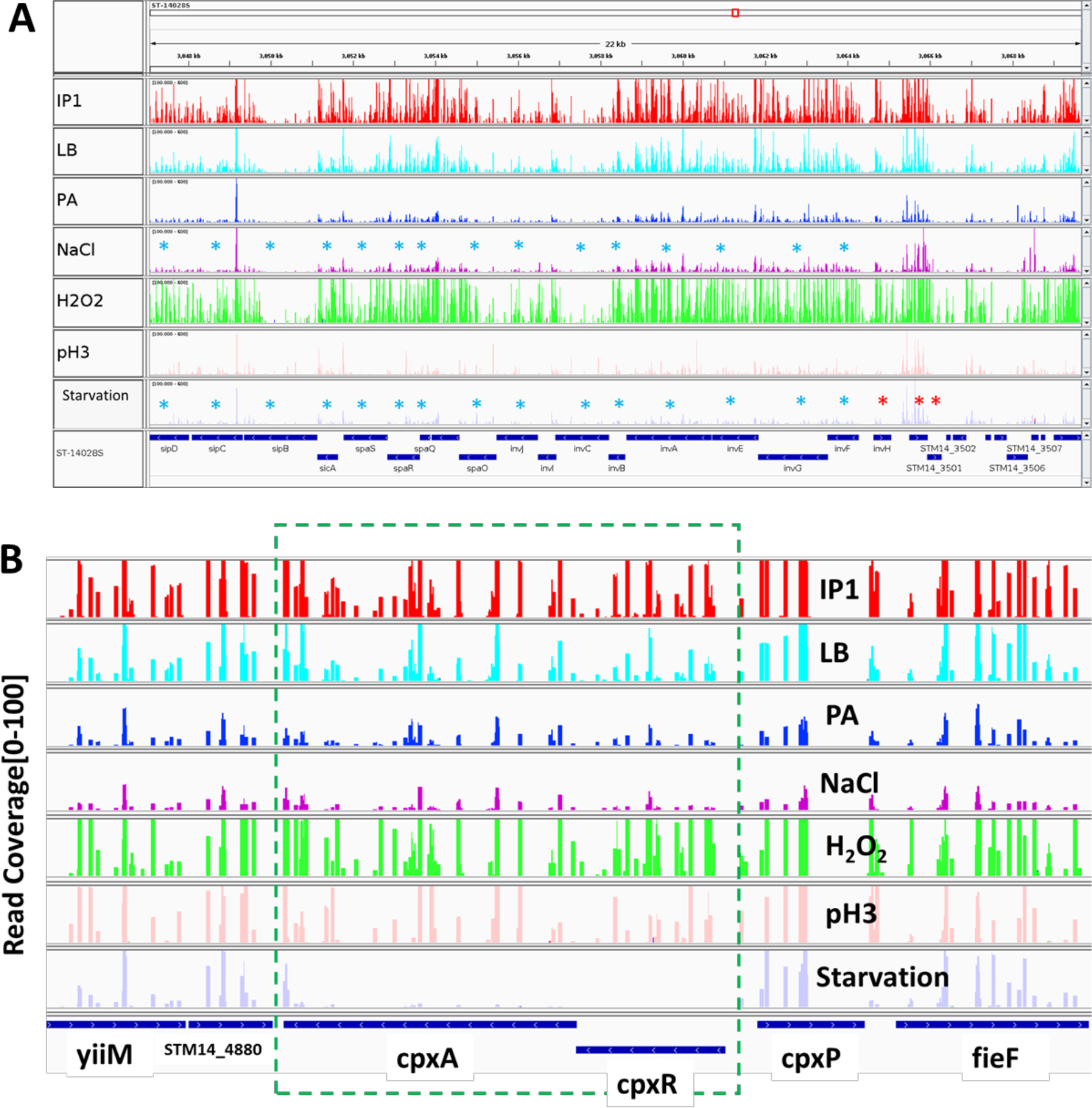
Tn-seq profiles around the selected genomic regions. A) *Salmonella* pathogenicity island 1 (SPI-1) genes encoding type III secretion system (TTSS). Screen shot image produced using Integrative Genomics Viewer (IGV) showing raw read coverage [100-600] in seven conditions. (Blue asterisk: conditionally essential in NaCl and Starvation; and Red asterisk: conditionally essential in Starvation only). B) CpxAR were conditionally essential in starvation. only.

**Figure S4.**
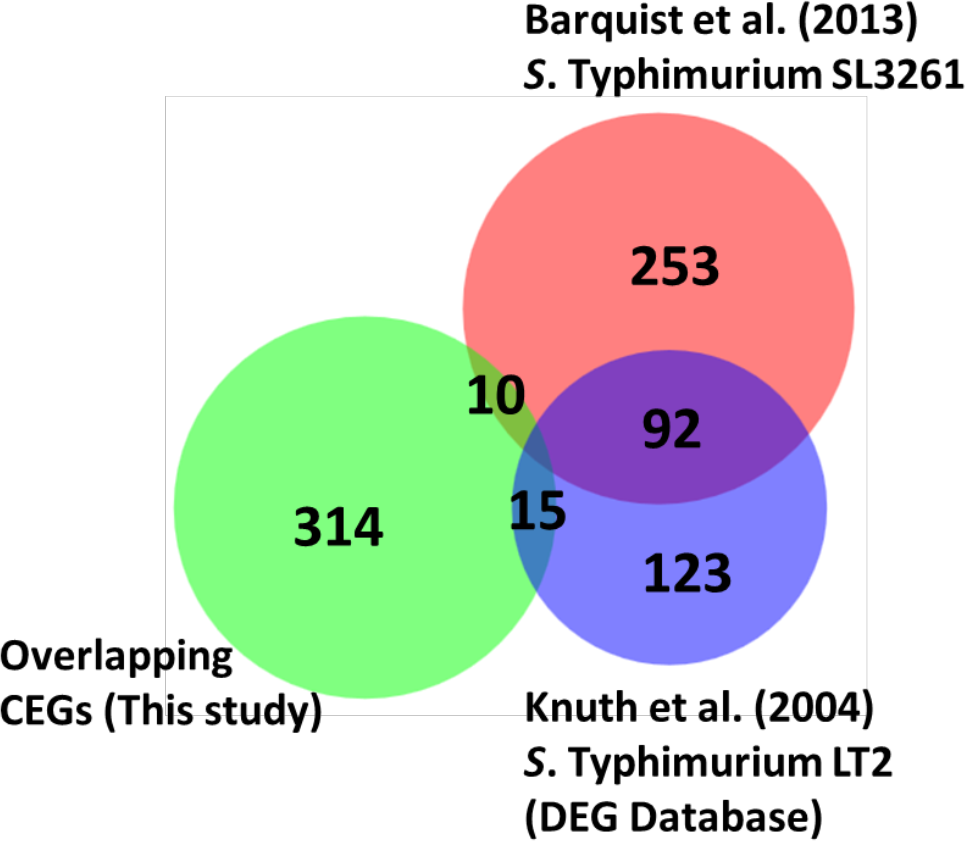
Comparison of the overlapping set of conditionally essential genes of *S*. Typhimurium 14028s (this study) with essential genome of *S*. Typhimurium SL3261 and *S*. Typhimurium LT2.

**Figure S5.**
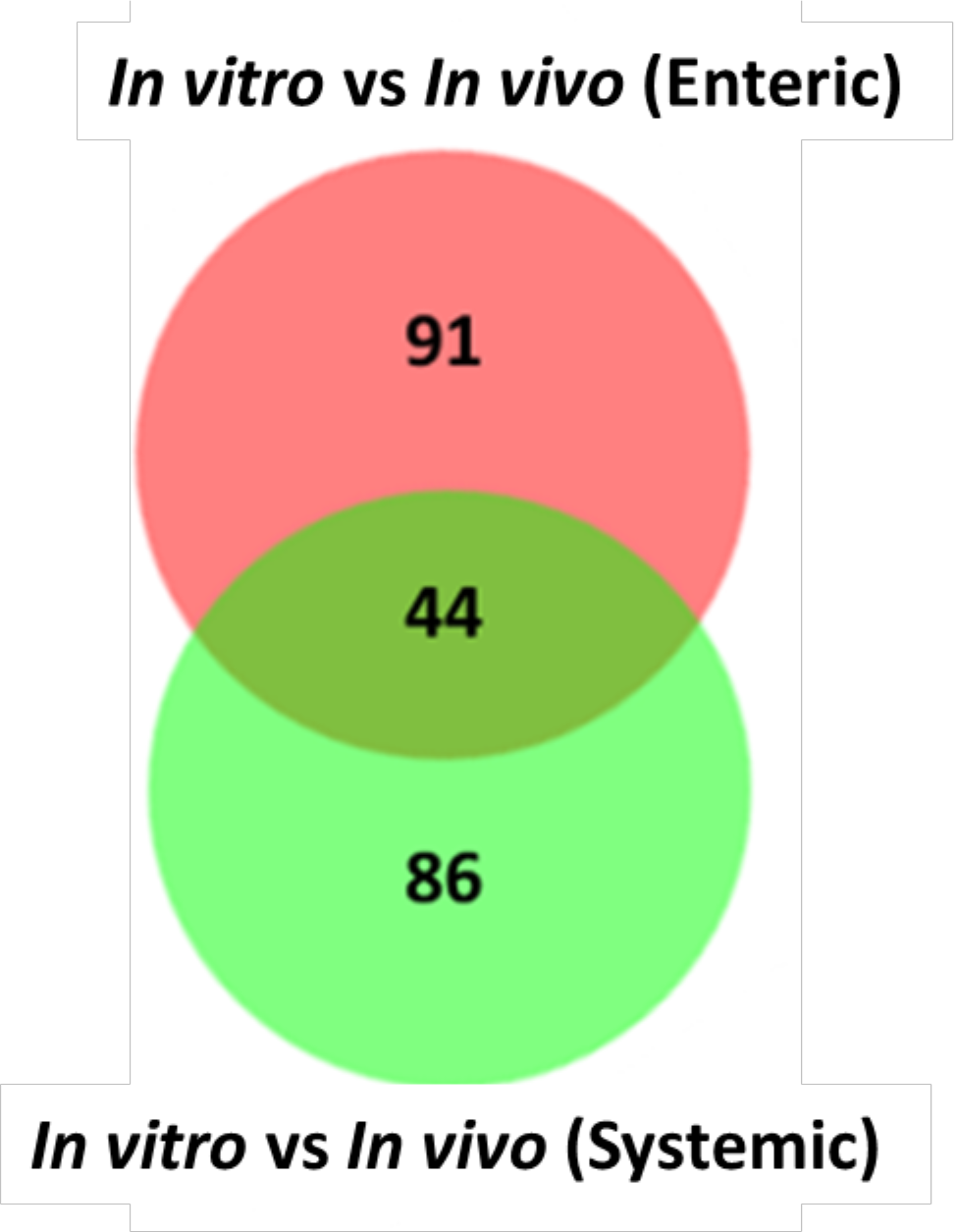
Genes required for enteric, systemic and *in vitro* fitness. Venn diagram shows the number of shared genes between *in vitro* vs *in vivo* (Enteric) (135 CEGs shown in Fig 5) and *in vitro* vs *in vivo* (Systemic) (130 CEGs shown Fig 6). The list of 44 genes required for *in vitro*, enteric and systemic fitness are shown in Table 1.

## List of Supplementary Tables

**Table S1.** Oligonucleotides used in this study.

**Table S2.** All conditionally essential genes (CEGs) in S. Typhimurium 14028S identified in this study.

**Table S3.** Comparison of the conditionally essential genes (CEGs) in S. Typhimurium 14028s identified in this study across the 5 stress conditions.

**Table S4.** The conditionally essential genes (CEGs) in S. Typhimurium 14028s identified in this study that are located in Salmonella Pathogenicity Islands.

**Table S5.** Comparison of the conditionally essential genes (CEGs) of S. Typhimurium 14028s (this study) with the essential genes of S. Typhimurium identified from previous studies

**Table S6.** The conditionally essential genes (CEGs) in the presence of the in vitro host stressors (PA, NaCl, pH3, Bile, and LB42) that are also required for enteric infection in farm animals (cattle, pig, and chicken).

**Table S7.** The conditionally essential genes (CEGs) in the presence of the in vitro host stressors (H2O2, NaCl, pH3, Starvation, and dLB) that are also required for systemic infection (MΦ; infections (MΦ, A-Mice, P-Mice, Sp-Liv).

